# TOM-1/Tomosyn acts with the UNC-6/Netrin receptor UNC-5 to inhibit growth cone protrusion in *Caenorhabditis elegans*

**DOI:** 10.1101/2022.06.16.496459

**Authors:** Snehal S. Mahadik, Erik A. Lundquist

**Affiliations:** Program in Molecular, Cellular, and Developmental Biology, Department of Molecular Biosciences, The University of Kansas, 1200 Sunnyside Avenue, 5049 Haworth Hall, Lawrence, KS 66045

## Abstract

In the polarity/protrusion model of growth cone repulsion from UNC-6/Netrin, UNC-6 first polarizes the VD growth cone via the UNC-5 receptor, and then regulates protrusion asymmetrically across the growth cone based on this polarity. Through the UNC-40/DCC receptor, UNC-6 stimulates protrusion dorsally, and through UNC-5 inhibits protrusion ventrally and laterally, resulting in net dorsal growth. Previous studies showed that UNC-5 inhibits growth cone protrusion via the flavin monooxygenases and potential destabilization of F-actin, and via UNC-33/CRMP and restriction of microtubule + end entry into the growth cone. To explore the role of vesicle fusion in growth cone protrusion, we analyzed *tom-1/tomosyn* mutants. Tomosyn normally occludes formation of the SNARE complex by interacting with and inhibiting Syntaxin and thus preventing vesicle fusion. VD growth cones of *tom-1 null* mutants were similar to wild-type. However, *tom-1* null mutants suppressed the effects of constitutively-activated MYR::UNC-5, which alone causes small growth cones with little protrusion. This suggests that TOM-1 is normally required for the inhibitory effects of MYR::UNC-5 on growth cone protrusion. *tom-1* encodes long and short isoforms, and results here indicate that *tom-1S* is required downstream of UNC-5 to inhibit protrusion, whereas the *tom-1L* has a pro-protrusive role. *unc-64/Syntaxin* mutants displayed reduced growth cone protrusion, suggesting that TOM-1 inhibits growth cone protrusion by inhibiting UNC-64/Syntaxin, similar to its role in neurotransmission. TOM-1L, TOM-1S, and UNC-64/Syntaxin were all required for VD growth cone polarity of protrusion, indicating that regulated vesicle fusion is required for the establishment and/or maintenance of VD growth cone polarity. These studies show that, in addition to effects on actin and microtubules, UNC-5 might inhibit VD growth cone protrusion by inhibiting growth cone vesicle fusion and thus the ability of growth cones to add plasma membrane necessary for protrusive growth.

## Introduction

The guided extensions of axons and dendrites play a crucial role in the development of neural circuit and networks. Growth cones present at the growing tip of the neurite guide the outgrowth of the neurite. Growth cones are dynamic structures that consist of a branched actin lamellipodial body and filopodial consisting of F-actin bundles (Gallo and Letourneau 1999; Gallo and Letourneau 2004; Zhou and Cohan 2004; Pak *et al*. 2008; Lowery and Van Vactor 2009). Extracellular guidance cues are detected by growth cones via receptors present on the plasma membrane which orchestrate a series of intracellular events that guide the growth cone in the proper direction (Tessier-Lavigne and Goodman 1996; Gallo and Letourneau 1999; Gallo and Letourneau 2004; Lowery and Van Vactor 2009), including actin cytoskeletal dynamics, microtubule delivery of vesicles and cytoskeletal regulators, and addition of plasma membrane via exocytosis.

In *C. elegans* and vertebrates, UNC-6/Netrin is a bifunctional, conserved, secreted laminin like guidance cue, which directs the dorsal-ventral axon guidance by utilizing its receptors UNC-40/DCC and UNC-5 (Hedgecock *et al*. 1990; Ishii *et al*.1992; Chan *et al*. 1996; Norris and Lundquist 2011). The VD motor neuron cell bodies reside in the ventral nerve cord and send processes anteriorly in the VNC, which then turn dorsally and migrate to the dorsal nerve cord, forming an axon commissure. The dorsal commissural growth of the VD growth cone is dependent upon UNC-6/Netrin in the ventral nerve cord (i.e. VD growth cones migrate away from UNC-6/Netrin, classically called repulsion) (Wadsworth *et al*. 1996; Wadsworth 2002; Norris and Lundquist 2011). As the VD growth cone migrates dorsally, it extends dynamic filopodial protrusions biased to the dorsal direction of growth (Knobel *et al*. 1999; Norris and Lundquist 2011).

Classically, it was thought that UNC-6/Netrin forms a ventral-to-dorsal gradient that growth cones dynamically sense (Tessier-Lavigne and Goodman 1996; Boyer and Gupton 2018). Growth up (attraction) or down (repulsion) the gradient involved the UNC-40/DCC receptor and UNC-5 respectively. Recent studies in vertebrate spinal cord suggest that gradients are not involved and that Netrin1 acts in a short-range,haptotactic mechanism in spinal cord commissural guidance (Dominici *et al*. 2017; Varadarajan and Butler 2017; Yamauchi *et al*. 2017; Morales 2018).

In *C. elegans, in vivo* imaging studies of VD growth cones in wild-type and *unc-6* signaling mutants also do not support the gradient model. Instead, they indicate that UNC-6/Netrin first polarizes the growth cone via the UNC-5 receptor, and then regulates growth cone protrusion based upon this polarity (Norris and Lundquist 2011; Norris *et al*. 2014; Gujar *et al*. 2018). UNC-5 inhibits protrusion ventrally, and UNC-40 stimulates protrusion dorsally, resulting in net dorsal growth. Thus, both UNC-5 and UNC-40 receptors act in the same growth cone in growth away from UNC-6/Netrin by balancing protrusion across the growth cone. This polarity/protrusion model is a new paradigm in which to understand UNC-6/Netrin in axon guidance. The statistically oriented asymmetric localization (SOAL) model involving UNC-40 acts similarly in growth cone growth toward UNC-6/Netrin (Kulkarni *et al*. 2013; Yang *et al*. 2014; Limerick *et al*. 2017).

Central to the polarity/protrusion model is that finding that UNC-5 polarizes the growth cone and then inhibits growth cone lamellipodial and filopodial protrusion based on this polarity. UNC-40 stimulates protrusion at the dorsal growing tip, and UNC-5 inhibits protrusion ventrally and laterally (Norris and Lundquist 2011; Norris *et al*. 2014; Gujar *et al*. 2018). Previous studies indicate that UNC-5 inhibits VD growth cone protrusion through two mechanisms. The flavin monooxygenases (FMOs) act downstream of UNC-5 to inhibit protrusion, possible by destabilizing F-actin similar to the FMO protein MICAL (Gujar *et al*. 2017). UNC-33/CRMP acts downstream of UNC-5 and inhibits protrusion by restricting microtubule + end entry into the growth cone (Gujar *et al*. 2018). Microtubules are pro-protrusive in the VD growth cones (Gujar *et al*. 2018; Gujar *et al*. 2019), possibly by delivering vesicles and cytoskeletal regulators such as Arp2/3 and Enabled (Norris *et al*. 2009).

Here a potential third pathway downstream of UNC-5 in inhibiting protrusion is explored, involving regulation of vesicle exocytosis and addition of plasma membrane to the growth cone. Vesicle exocytosis is required for growth cone membrane extensions by fusion of plasmalemmal precursor vesicles at the plasma membrane of growth cones and is closely tied to cytoskeletal dynamics (Futerman and Banker 1996; Hausott and Klimaschewski 2016; Nozumi and Igarashi 2018), and a balance of endocytosis and exocytosis is involved in growth cone guidance (Tojima *et al*. 2014; Tojima and Kamiguchi 2015). The axon guidance cue reelin controls fusion of VAMP7-positive vesicles in regenerating dorsal root ganglion neurons (Jausoro and Marzolo 2021). Furthermore, synaptic-like vesicles in the growth cone are required for pioneer axon navigation in zebrafish (Nichols and Smith 2019). In *C. elegans*, the RAB-3 GTP exchange factor AEX-3 controls pioneer axon guidance (Bhat and Hutter 2016).Studies on rat cultured hippocampal neurons suggest that exocytosis is restricted to the distal, dynamic region of growth cones which leads to the membrane addition and extension (Sakisaka *et al*. 2004).

Tomosyn was identified as a Syntaxin-interacting molecule that was shown to inhibit interaction of the T-SNARE Syntaxin with the V-SNARE Synaptobrevin, thus blocking formation of the SNARE complex and preventing vesicle fusion (Fujita *et al*. 1998). Tomosyn has been well-characterized in inhibition of synaptic vesicle fusion and dense core vesicle fusion in the nervous system (Fujita *et al*. 1998; Hatsuzawa *et al*. 2003; Widberg *et al*. 2003; Pobbati *et al*. 2004; McEwen *et al*. 2006; Takamori *et al*.2006; Gladycheva *et al*. 2007), including *C. elegans* (Dybbs *et al*. 2005; Gracheva *et al*. 2006; McEwen *et al*. 2006; Gracheva *et al*. 2010; Burdina *et al*. 2011). Tomosyn also regulates growth cone morphology in cultured rat hippocampal neurons. Vesicle fusion is inhibited in the proximal “palm” of the growth cone due to the action of Tomosyn in this region (Sakisaka *et al*. 2004). When growth cones undergo collapse, Tomosyn relocalizes around the perimeter of the growth cone (Sakisaka *et al*. 2004).

In this work, the role of TOM-1 is explored in VD growth cone morphology and interaction with UNC-5 in inhibiting VD growth cone protrusion. While complete loss of TOM-1 had little effect on VD growth cone morphology, it did suppress growth cone inhibition of protrusion driven by activated MYR::UNC-5. This suggests that TOM-1 is required for MYR::UNC-5 to inhibit growth cone protrusion and that TOM-1 might act downstream of UNC-5 in this process. *tom-1* encodes long and short isoforms. The long isoforms contain the N-terminal WD40 repeats and the C-terminal R-SNARE domain that interacts with Syntaxin, and the short isoform only encodes the C-terminal R-SNARE domain and lacks the WD40 repeats. A mutation that eliminates only the long isoforms but not the short isoform resulted in small, less-protrusive VD growth cones. Furthermore, *tom-1L* mutation suppressed the excess VD growth cone protrusion observed in *unc-5* loss-of-function mutants. These results suggest that the TOM-1L and TOM-1S isoforms have opposing functions, with TOM-1S acting in an anti-protrusive manner and TOM-1L acting pro-protrusive. Mutation of *tom-1S* without affecting *tom-1L* resembled the complete loss of *tom-1* (no effect on VD growth cone alone but suppressed *myr::unc-5*). Furthermore, transgenic expression of *tom-1S* resulted in reduced VD growth cone protrusion. Taken together, these data suggest that in VD growth cones, *tom-1S* is the active isoform in inhibiting growth cone protrusion and possibly vesicle fusion, whereas *tom-1L* has a pro-protrusive role. Finally, a hypomorphic *unc-64/Syntaxin* mutant had small, less protrusive VD growth cones and suppressed the excess protrusion of *unc-5* loss-of-function mutants. In sum, these results are consistent with a model wherein UNC-5 engages TOM-1S to inhibit vesicle fusion and thus inhibit growth cone protrusion. Additionally, *tom-1L, tom-1S*, and *unc-64/Syntaxin* were each required for VD growth cone polarity of protrusion, suggesting that regulated vesicle fusion is necessary to establish and/or maintain VD growth cone polarity.

## Results

### *tom-1* encodes long and short isoforms

The *C. elegans* genome encode a single *Tomosyn* gene, *tom-1*, which was identified in a forward genetic screen for enhancers of acetylcholine secretion (Dybbs *et al*. 2005). TOM-1 is an ortholog of mammalian Tomosyn and shares a significant sequence similarity with mammalian Tomosyn-1 and Tomosyn-2 (Gracheva *et al*. 2006). Through alternative 5’ end usage and alternative splicing, *tom-1* produces multiple isoforms. Long isoforms include *tom-1A* (Figure 1A). A short isoform is encoded by *tom-1B* (Figure 1A), produced by alternative 5’ exon located in an intron of *tom-1A*. RNA-seq on three independent replicates of mixed-stage wild-type animals revealed that the *tom-1B* 5’ exon and splice occurred in 8/137, 9/151 and 4/109, or ∼5%, of splicing events involving this intron (Figure 1B). This is likely to be an underestimate due to the location of this small exon at the 5’ end on the transcript and not being completely represented during RNA-seq library construction. Long isoforms of *tom-1* encode conserved WD40 repeats in the N-terminus, and a conserved R-SNARE-like domain at the C-terminus, through which Tomosyn interacts with the SNARE complex (Figure 1A) (Gracheva *et al*. 2006). The short *tom-1B* isoform lacks the N-terminal WD40 repeats and encodes only the C-terminal R-SNARE domain.

**Figure 1.**
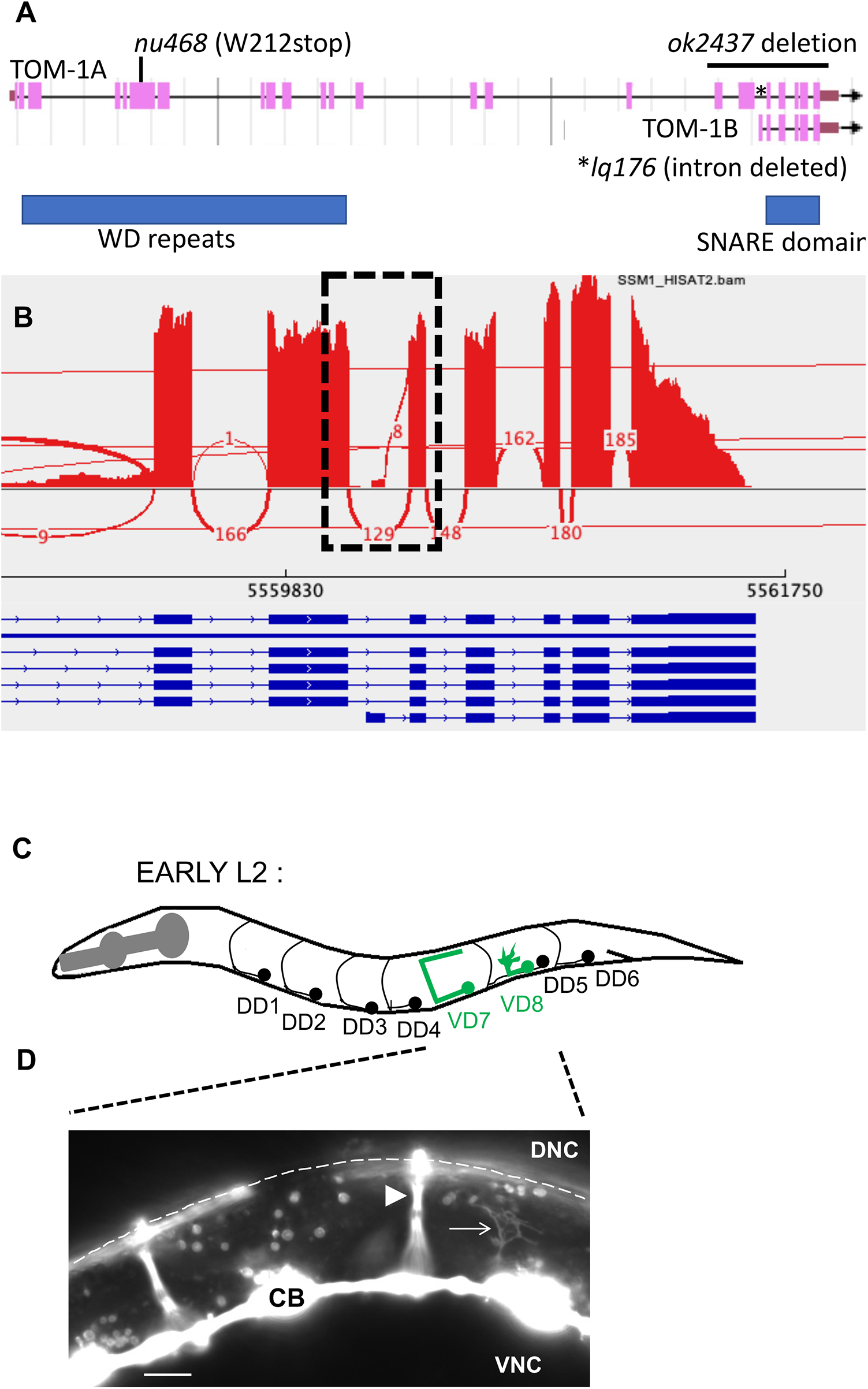
*tom-1* gene structure and VD growth cones. (A) The *tom-1* gene structure is shown (from Wormbase WS284). *tom-1A* is representative of the long isoforms, and *tom-1B* is the short isoform. Regions encoding the WD40 repeats, and the SNARE domain are indicated by blue boxes. Alleles and location of their molecular lesions producing multiple isoforms. *tom-1(nu468)* results in a premature stop codon predicted to affect all long isoforms but not *tom-1B* short. *tom-1(ok2437*) is a 2391-bp deletion that removes all of exons 17 to 23 and is predicted to affect all isoforms of *tom-1. tom-1(lq176)* is a precise deletion of intron 18 of *tom-1A*, leaving the coding potential for *tom-1A* unchanged but deleting the first exon of *tom-1B*. (B) A Sashimi plot from the Integrated Genome Viewer of splice junctions in RNA-seq reads of *tom-1*. RNA-seq was conducted on mixed stage N2 animals, reads aligned using HISAT2, and analyzed with the Integrated Genome Viewer. Gene structure is below in blue, and splice junctions with their abundance in the RNA-seq reads is indicated. The dashed box shows splices at intron 18, of which 8/137 include the *tom-1B* first exon. (C) Diagram of an early L2 larva of *C. elegans* hermaphrodite highlighting the structure and position of the DD motor neurons and axons (black), and VD7 and VD8 (green). Dorsal is up, and anterior is left. A growth cone is represented on the VD8 axon as it extends dorsally. (D) A fluorescent micrograph of an early L2 animals with *unc-25::gfp* expression in the VD and DD neurons. Dorsal is up, and anterior is left. An arrowhead indicates a DD commissural axon, and an arrow points to a VD growth cone as it extends dorsally. CB, cell body; DNC, dorsal nerve cord; VNC, ventral nerve cord. Scale bar represents 5μm.

*tom-1(ok2437)* is a 2391 bp deletion that removed 3’ exons predicted to affect all isoforms of *tom-1* (Figure 1A), and *tom-1(nu468)* introduces a premature stop codon at tryptophan at 212 (Dybbs *et al*. 2005) and is predicted to affect only the long isoforms and not the *tom-1B* short isoform (Figure 1A).

### TOM-1 regulates VD growth cone protrusion and polarity

In the early L2 larval stage, the axons of the VD neurons begin their ventral-to-dorsal commissural growth, with a visible growth cone at the tip of the extending commissural VD axons (Figure 1C and D). DD axons extend earlier, in late embryogenesis. *tom-1(nu468)* and *tom-1(ok2437)* mutants both displayed low levels of VD/DD axon guidance defects (Figure 2 B, C, D; Figure 3A).

**Figure 2.**
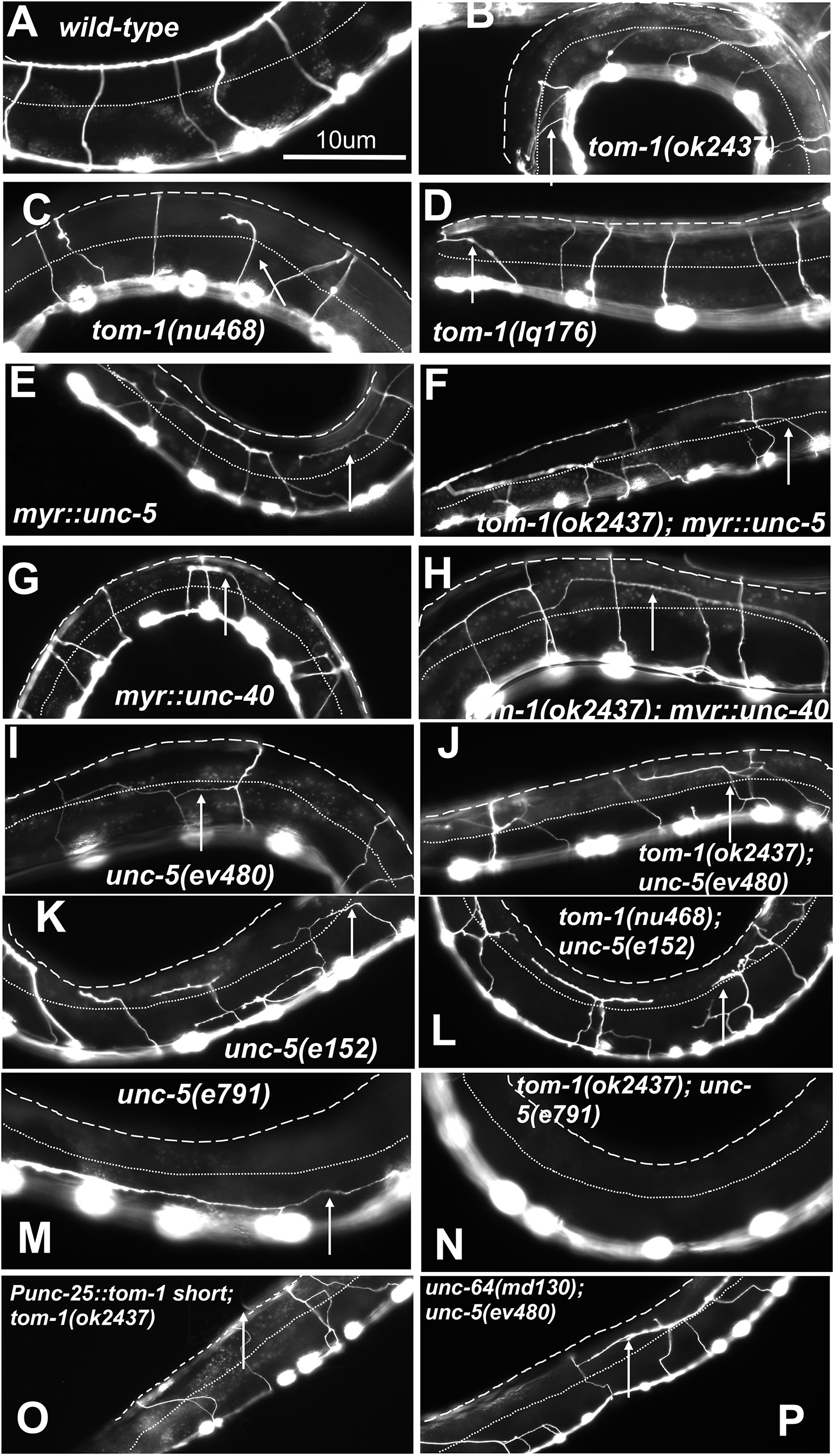
VD/DD axon guidance defects. Shown are fluorescent micrographs of the *Punc-25::gfp* transgene *juIs76* expressed in the VD/DD neurons of L4 animals. Dorsal is up and anterior left. The approximate lateral midline is indicated with a dotted white line, and the dorsal nerve cord by a dashed white line. White arrows indicate axon guidance defects in each genotype. Scale bar in (A) represents 10μm. Genotypes are indicated in each figure panel.

**Fig 3.**
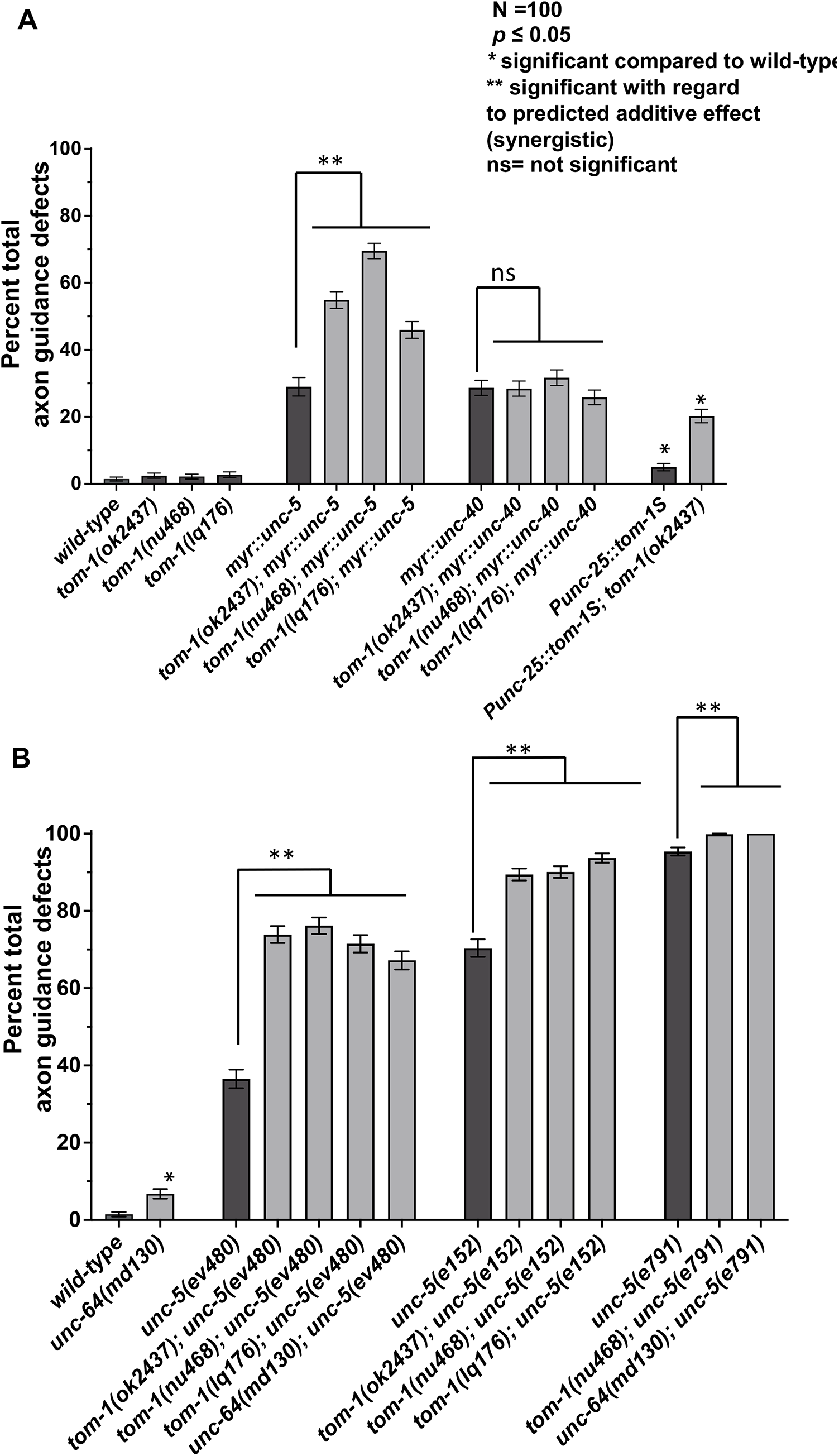
Quantification of VD/DD axon guidance defects. Total axon guidance defects were quantified as described in Materials and Methods. Genotypes are listed on the X axis, and total axon guidance defects on the Y axis. Error bars represents 2x standard error of proportion. Significance of difference was determined using Fisher’s exact test. Single asterisks (*) indicate significance compared to wild-type; double asterisks (**) indicate significance of single mutants alone to the predicted additive effect of double mutants calculated by the formula p1 + p2 – (p1p2). ns= not significant.

Growth cone morphology of VD axons in *tom-1* mutants was analyzed as previously described (Norris and Lundquist 2011; Norris *et al*. 2014; Mahadik and Lundquist 2022). Wild-type VD growth cones displayed an average area of 4.6μm^2^(Figure 4A and D), and filopodial protrusions with an average length of 0.9μm (Figure 4B and D). VD growth cones are polarized, with filopodial protrusions biased to the dorsal aspect of the growth cone, the direction of growth (Figure 4C and D). *tom-1(ok2437)* null mutants displayed a slight but not statistically significant increase in VD growth cone area and filopodial length (Figure 4 A, B, and E). The long isoform-specific *tom-1(nu468)* mutant VD growth cones displayed significantly reduced growth cone area and filopodial length (Figure 4 A, B, and F). These results suggest that *tom-1* long isoforms might have a pro-protrusive role in the growth cone.

**Figure 4.**
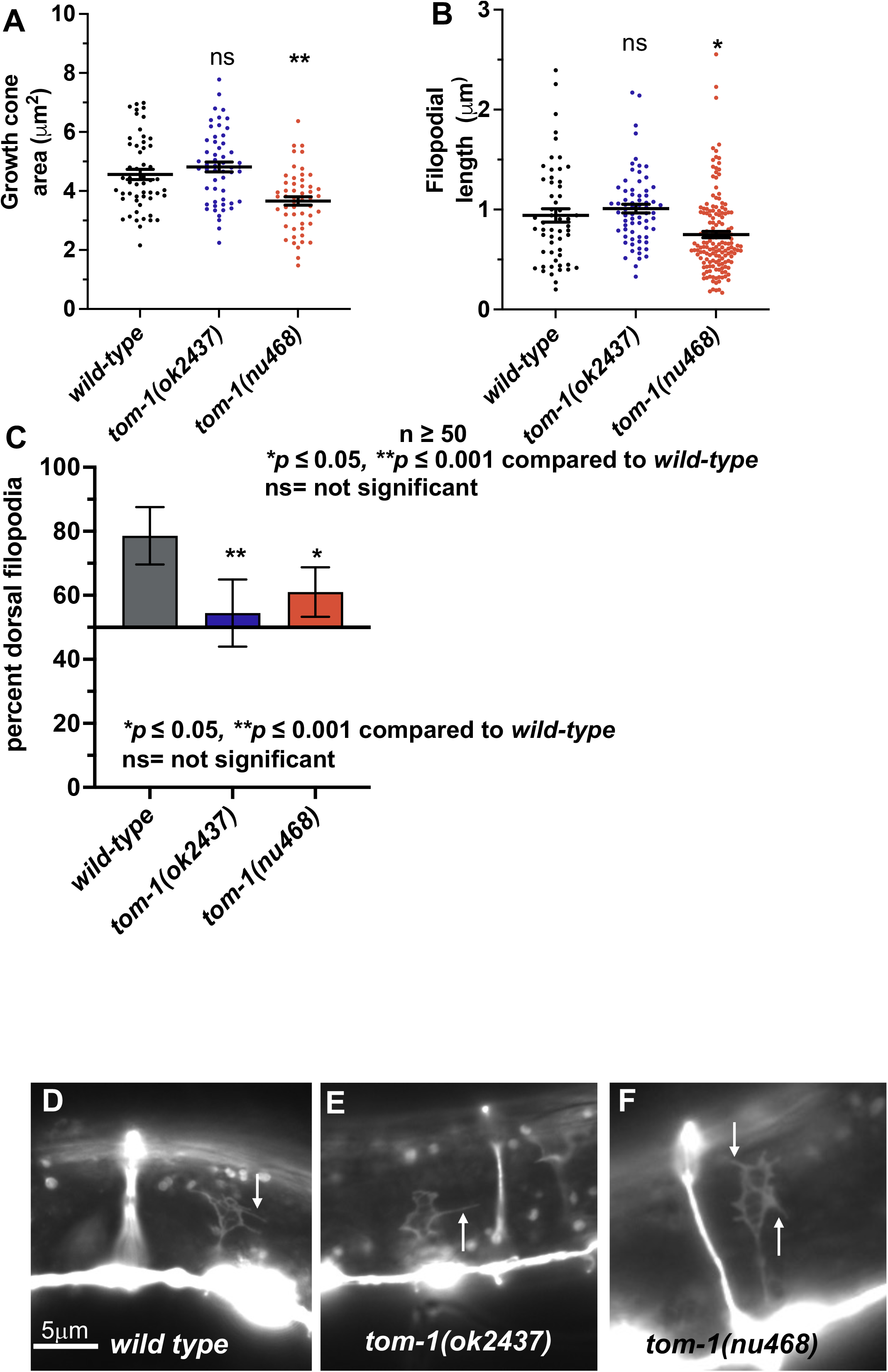
Growth cone analysis in *tom-1* mutants. At least 50 growth cones were scored in each genotype. (A, B) Quantification of VD growth cone area and filopodial length (see Materials and Methods). In the graph, each point represents a measurement of a single growth cone or filopodium. (A) Area of growth cone, in μm^2^. (B) Filopodial length, in μm. Error bars indicate standard error of the mean. Two-sided *t-*tests with unequal variance were used to determine the significance between wild-type and mutants. n = number of growth cones. Single asterisks (*) indicate significance at *p* < 0.05. Double asterisks (**) indicate significance at *p* < 0.001. C) A graph showing the percent of dorsally directed filopodial protrusions in VD growth cones of different genotypes (see Materials and Methods). The X-axis is set at 50%, such that bars extending above the X-axis represents above 50%, and bars that extends below represents below 50%. In wild-type, a majority of filopodia (78%) extended from the dorsal half of the growth cone. Significance between wild-type and mutants was determined by Fisher’s exact test. Error bar represents 2x standard error of proportion. n = number of growth cones. Single asterisks (*) indicates the significant *p* < 0.05 double asterisks (**) indicate significant *p* < 0.001. (D-F) Fluorescence micrographs of wild-type and mutant VD growth cones expressing *Punc-25::gfp*. (D, E, F). Arrows point to filopodial protrusions. Dorsal is up; anterior is left. The scale bar in (D) represents 5μm.

Dorsal polarity of growth cone protrusion was significantly reduced in both *tom-1(ok2437)* and *tom-1(nu468)* mutants (Figure 4C-F). These data suggest that both long and short TOM-1 isoforms are required for growth cone polarity of filopodial protrusion. That the long isoform-specific *tom-1(nu468)* mutants displayed reduced growth cone area and shorter filopodial protrusions and the null *tom-1(ok2437)* mutants did not suggest that the long and short isoforms of TOM-1 might have distinct roles in regulation of VD growth cone morphology.

### TOM-1 short is required for the effects of activated MYR::UNC-5 on VD growth cone morphology

Previous work showed that UNC-6/Netrin signaling regulates VD growth cone protrusion, including growth cone area, filopodial length, and polarity (Norris and Lundquist 2011; Norris *et al*. 2014; Gujar *et al*. 2017). The UNC-6/Netrin receptor UNC-5 normally inhibits growth cone protrusion. Constitutively-active MYR::UNC-5 expression in VD growth cones results in small growth cones with few and shortened filopodial protrusions (Figure 5A, B, and D). UNC-5 can act as a heterodimer with the UNC-6/Netrin receptor UNC-40, and constitutively-activated MYR::UNC-40 also results in reduced growth cone protrusion (Figure 5A, B, and G) (Gitai *et al*. 2003; Norris and Lundquist 2011; Norris *et al*. 2014; Gujar *et al*. 2017).

**Figure 5.**
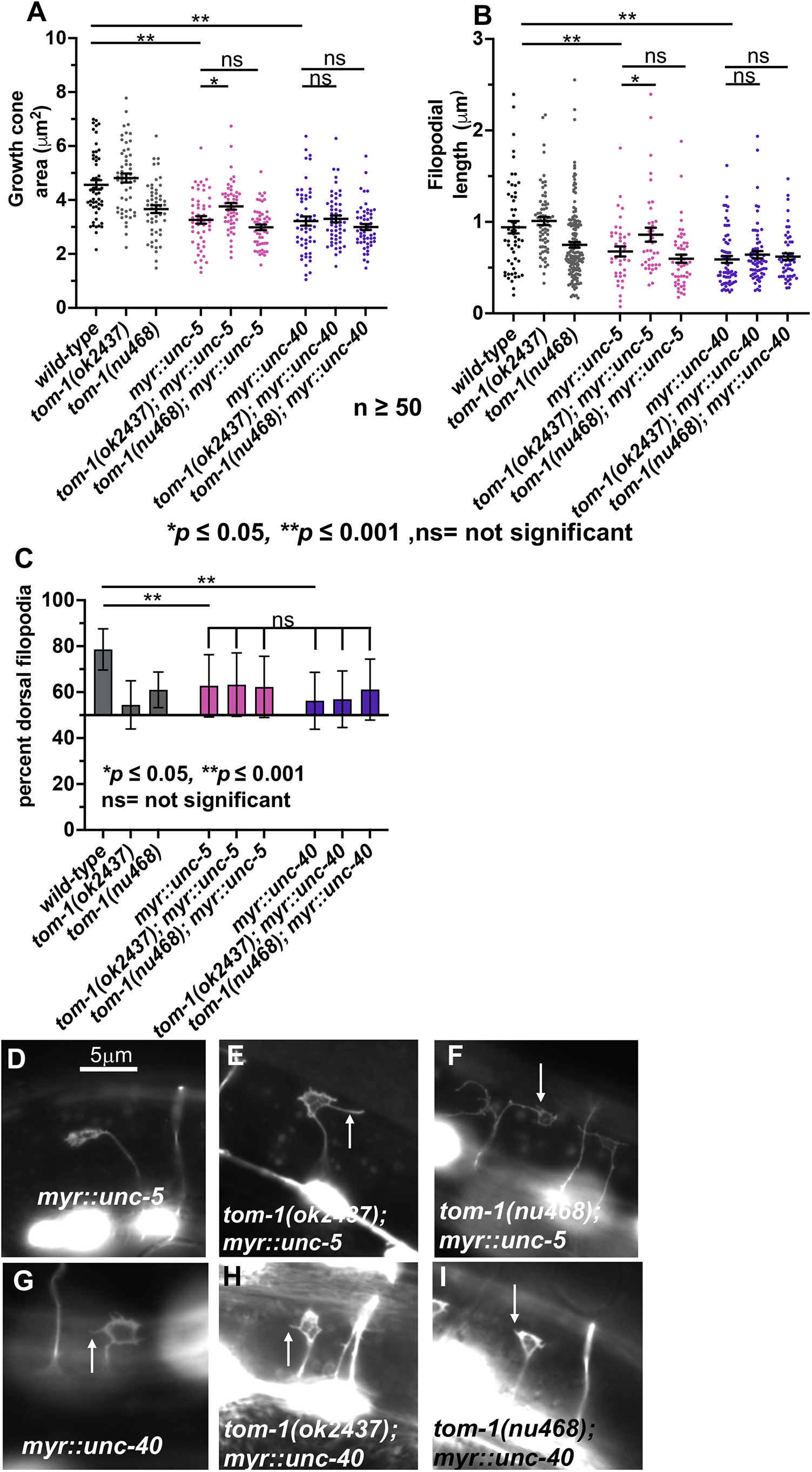
Growth cone analysis of *tom-1* with *myr::unc-5* and *myr::unc-40*. A, B) Quantification of VD growth cone area and filopodial length as described in Figure 4. Measures of significance shown on previous figures are not indicated here. The lines indicate genotypes analyzed for significance. C) A graph showing the percent of dorsally-directed filopodial protrusions in VD growth cones of different genotypes as described in Figure 4. *myr::unc-5* and *myr::unc-40* significantly perturbed growth cone polarity (lines), but no *tom-1* double mutant showed any significant difference (brackets). (D-I) Fluorescence micrographs of mutant VD growth cones of indicated genotypes expressing *Punc-25::gfp*. Arrows point to filopodial protrusions. Dorsal is up; anterior is left. The scale bar in (D) represents 5μm.

The *tom-1(ok2437)* null mutation significantly suppressed the effects of *myr::unc-5* (Figure 5A, B, and E). *tom-1(ok2437); myr::unc-5* growth cones were larger with longer filopodia compared to *myr::unc-5* alone. In contrast, *tom-1(ok2437)* had no effect on growth cone area or filopodial reduction caused by *myr::unc-40* (Figure 5A, B, and H). While loss of *tom-1* alone caused no VD growth cone phenotype, a role of TOM-1 in inhibiting growth cone protrusion was revealed in these studies using the *myr::unc-5* sensitized background. These results are consistent with TOM-1 acting downstream of UNC-5 to inhibit growth cone protrusion. As *myr::unc-40* was unaffected, TOM-1 might act specifically downstream of UNC-5 and not UNC-5::UNC-40 heterodimers.

### TOM-1 is required for inhibition of growth cone protrusion by MYR::UNC-5

*tom-1(nu468)*, which affects only the *tom-1* long isoforms and not the short *tom-1B* isoform, did not suppress the inhibition of growth cone area and filopodial length caused by *myr::unc-5* or *myr::unc-40* (Figure 5 A, B, F, and I). This suggests that the TOM-1 long isoforms are not required to inhibit growth cone protrusion. Alone, *tom-1(nu468)* mutants displayed reduced VD growth cone protrusion (Figure 4), suggesting a pro-protrusive role of the long isoforms. Thus, TOM-1 short and long isoform might have opposing roles, with TOM-1 short normally inhibiting protrusion and TOM-1 long normally stimulating protrusion. These data also suggests that the inhibitory function of short isoform of TOM-1 does not require long isoform.

Neither *tom-1(ok2437)* or *tom-1(nu468)* modified growth cone polarity of *myr::unc-5* or *myr::unc-40* (Figure 5C), consistent with both *tom-1* mutants alone affecting polarity of protrusion (Figure 4C).

*tom-1; myr::unc-5* double mutants showed a synergistic increase in the VD/DD axon guidance defects but *tom-1; myr::unc-40* double mutants did not (Figure 2, E, F, G, H) (Figure 3), consistent with TOM-1 specifically interacting with UNC-5. It is noted that despite restoration of MYR::UNC-5 growth cone protrusion by *tom-1(ok2437)*, axon guidance defects were increased. This suggests that growth cone protrusion is not the only aspect of growth cone dynamics during axon guidance affected by these molecules.

### TOM-1 long isoforms are required for excess growth cone protrusion in *unc-5* loss-of-function mutants

UNC-5 has been previously shown to inhibit VD growth cone protrusion (Norris and Lundquist 2011; Norris *et al*. 2014). As previously reported, *unc-5(e791)* strong loss-of-function mutants displayed increased growth cone area and filopodial length (Figure 6A, B, and G). *unc-5(ev480)* and *unc-5(e152)* are hypomorphic alleles that retain some UNC-5 function (Merz *et al*. 2001; Killeen *et al*. 2002; Mahadik and Lundquist 2022). However, *unc-5(ev480)* and *unc-5(e152)* displayed VD growth cone morphology similar to the *unc-5(e791)* null (Figure 6A, B, D, G). VD growth cones of *unc-5* mutants also lack the dorsal bias of growth cone protrusions (Figure 6C). *unc-5(e791)* mutants displayed a nearly complete failure of VD/DD axons to reach the dorsal nerve cord, *unc-5(ev480)* and *unc-5(e152)* displayed weaker axon guidance defects due to their hypomorphic nature (Figure 3).

**Figure 6.**
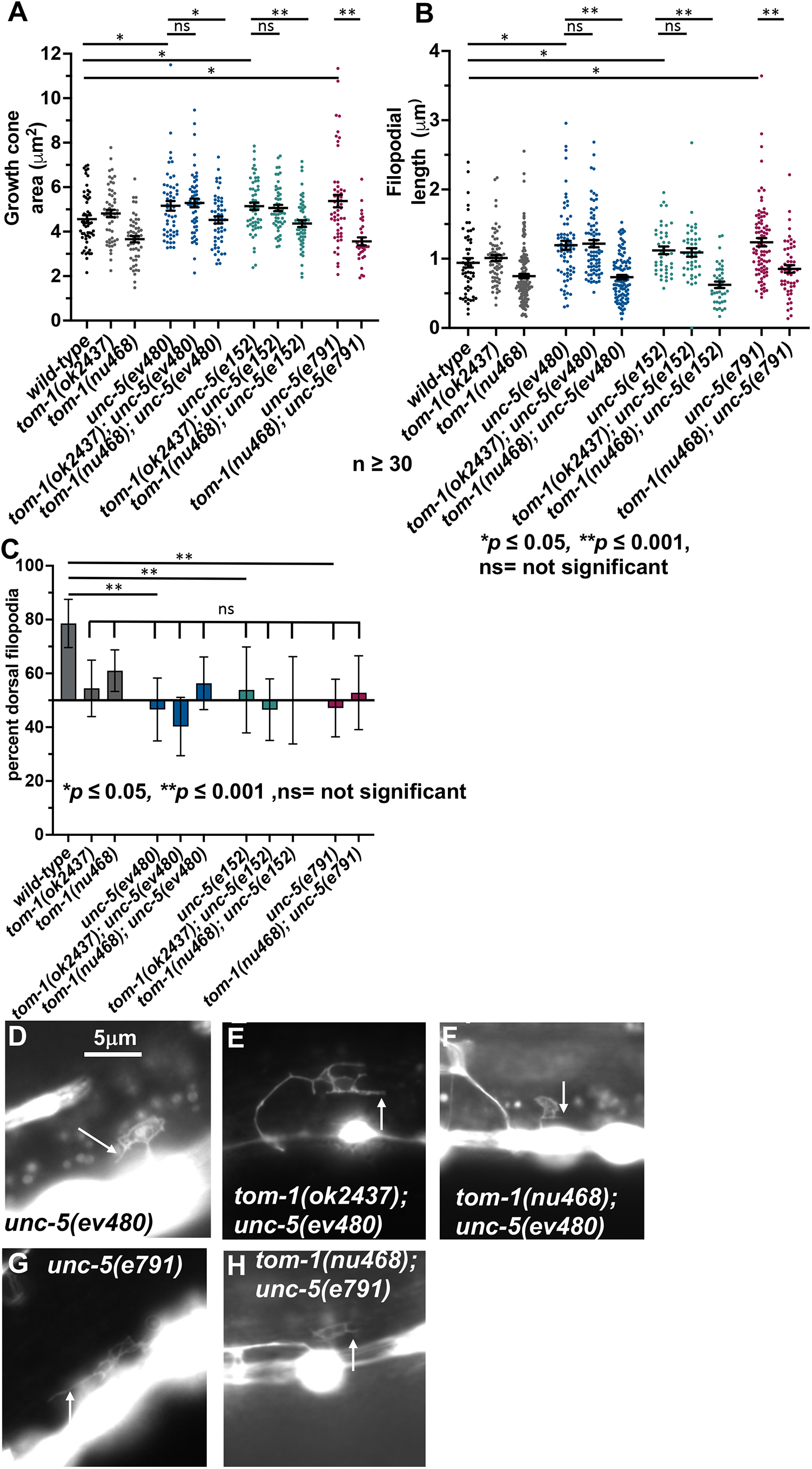
Growth cone analysis of *unc-5* and *tom-1* double mutants. A, B) Quantification of VD growth cone area and filopodial length as described in Figure 4. Measures of significance shown on previous figures are not indicated here. Lines indicate significance of comparisons between genotypes. C) A graph showing the percent of dorsally-directed filopodial protrusions as described in Figure 4. *unc-5* mutants significantly perturbed growth cone polarity (lines), but no *tom-1* double mutant showed any significant difference (brackets). (D-H) Fluorescence micrographs of mutant VD growth cones of indicated genotypes expressing *Punc-25::gfp*. Arrows point to filopodial protrusions. Dorsal is up; anterior is left. The scale bar in (D) represents 5μm.

VD growth cones of double mutants of the *tom-1(ok2437)* null allele and the *unc-5(e791)* null allele could not be scored because none emerged from the ventral nerve cord. There was a resulting complete failure of and VD or DD axons extending out of the ventral nerve cord (Figure 2 M and N). VD growth cones of *tom-1(ok2437)* with hypomorphic *unc-5(e152)* and *unc-5(ev480)* were apparent, and they resembled the growth cones of *unc-5* hypomorphs alone, with increased growth cone area and filopodial length compared to wild-type (Figure 6 A, B, E). Also, synergistic enhancement of VD/DD axon guidance defects compared to *unc-5(ev480, e152)* alone were observed (Figure 2 I and J, and Figure 3). Thus, *tom-1(ok2437)* did not alter the growth cone phenotypes of *unc-*5 hypomorphs, and in fact enhanced VD/DD axon guidance defects.

In contrast, the long isoform specific *tom-1(nu468)* mutant significantly suppressed growth cone area and filopodial length of all *unc-5* mutants (Figure 6 A, B, F, H). Despite suppressing excessive growth cone protrusion, there was a synergistic enhancement in VD/DD axon guidance defects of these double mutants (Figure 2 K, L) (Figure 3).

VD growth cones in all *tom-1; unc-5* mutants in which they were apparent were unpolarized similar to all of the single mutants alone (Figure 6C).

These studies indicate that the TOM-1 long isoforms were in part necessary for excessive growth cone protrusion in *unc-5* mutants, consistent with a pro-protrusive role of the TOM-1 long isoforms. This, combined with *tom-1* null suppression of activated MYR::UNC-5, suggest that the TOM-1 long and short isoforms have opposing roles in regulating growth cone protrusion, with TOM-1 long isoforms being pro-protrusive and TOM-1 short inhibiting protrusion.

### A *tom-1B* short isoform-specific mutant resembles the *tom-1* null

The *tom-1B* short isoform is produced by an alternative 5’ exon located in an intronic region of the long isoform (Figure 1A). We used Cas9 genome editing to precisely delete the intron contain the *tom-1B* 5’ exon, effectively fusing together exons 18 and 19 without affecting coding capacity of these exons (Figure 1). This *tom-1(lq176)* allele is predicted to encode all of the long isoforms.

*tom-1(lq176)* did not affect the growth cone area and filopodial length compared to wild-type (Figure 7 A, B, D, and E), but did display growth cone polarity defects (Figure 7C). *tom-1(lq176)* suppressed the inhibition of growth cone area and filopodial length caused by *myr::unc-5* (Figure 7 A, B, F, and G), but did not suppress the excessive growth cone protrusion hypomorphic *unc-5(ev480 and e152)* mutants (Figure 7 A, B, H, and I). No VD growth cones were apparent in *tom-1(lq176); unc-5(e791)* null double mutants. All double mutants with *tom-1(lq176), unc-5* hypomorphs, and *myr::unc-5* displayed slightly increased VD/DD axon guidance defects, in some cases significant (Figure 3). Furthermore, VD growth cone polarity in double mutants resembled each single mutant alone, which all had unpolarized growth cones (Figure 7C).

**Figure 7.**
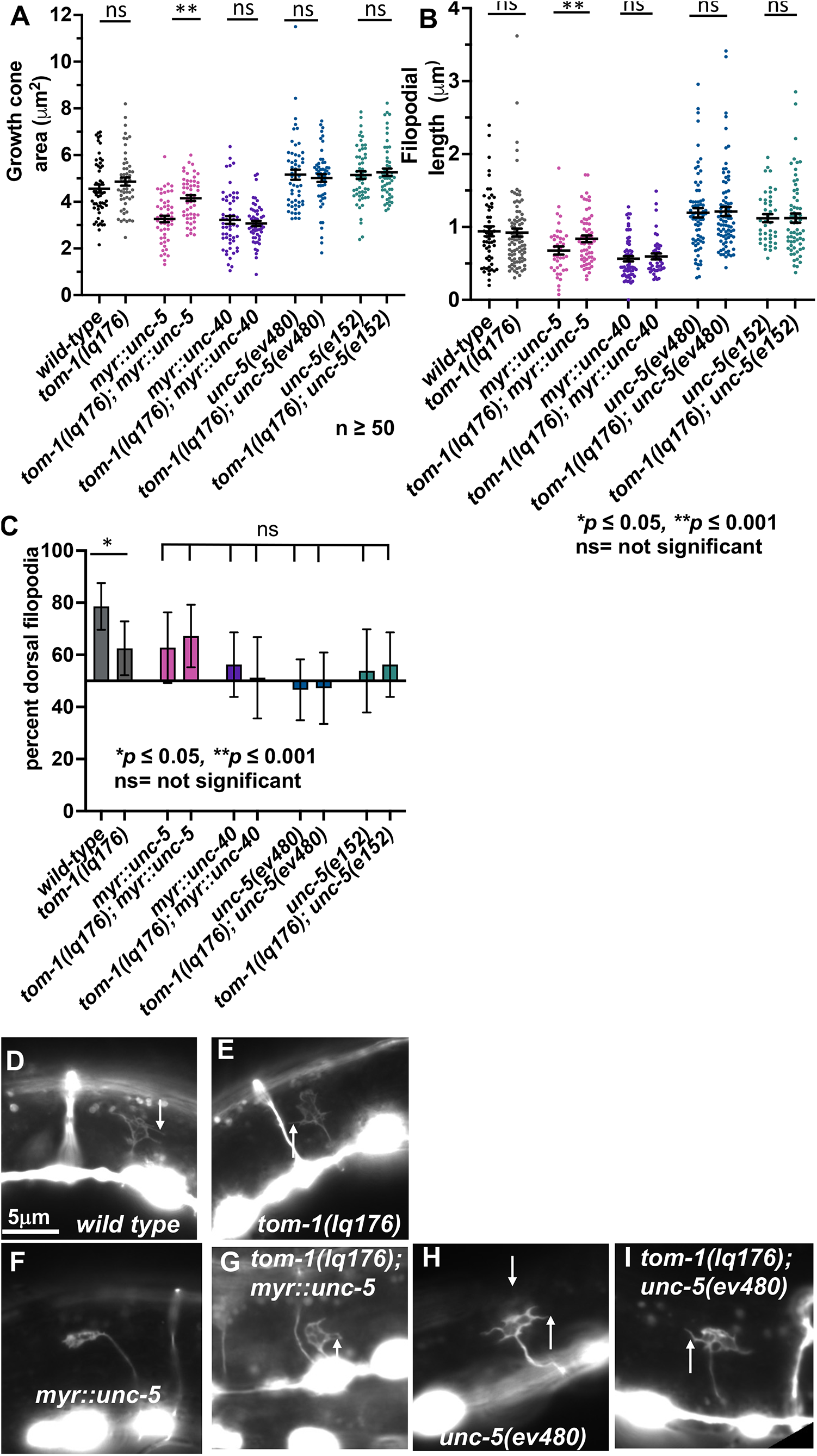
*tom-1(lq176)* growth cone quantification. (A, B) Quantification of VD growth cone area and filopodial length as described in Figure 4. Comparisons of significance indicated on previous figures are not shown here. Lines indicate significance of comparisons between genotypes. C) A graph showing the percent of dorsally-directed filopodial protrusions as described in Figure 4. *tom-1(lq176)* mutants significantly perturbed growth cone polarity (lines), but no *unc-5* double mutant showed any significant difference (brackets). (D-I) Fluorescence micrographs of mutant VD growth cones of indicated genotypes expressing *Punc-25::gfp*. Arrows point to filopodial protrusions. Dorsal is up; anterior is left. The scale bar in (D) represents 5μm.

In sum, *tom-1(lq176)* resembled the *tom-1(ok2437)* null mutant in all respects, including loss of growth cone polarity, suppression of growth cone inhibition by activated *myr::unc-5*, and failure to suppress excess growth cone protrusion in *unc-5* hypomorpic mutants. This indicates that the TOM-1 long isoforms alone cannot supply full TOM-1 function and is consistent with long and short isoforms having opposing roles (long are pro-protrusive; short is anti-protrusive).

### Transgenic expression of the TOM-1B short isoform inhibited VD growth cone protrusion

The genomic region of the *tom-1B* short isoform was placed under the control of the *unc-25* promoter expressed in the VD/DD neurons. Transgenic expression of this construct in a wild-type background resulted in reduced VD growth cone area and filopodial length (Figure 8 A, B, and F). Transgenic expression of *tom-1B* also abolished VD growth cone polarity of protrusion. Similar results were seen in the *tom-1(ok2437)* background (Figure 5 A, B, G). These data indicate that the TOM-1B short isoform inhibits growth cone protrusion, consistent with the loss of function studies of *tom-1(lq176)*. These data also indicate that *tom-1B* can act cell-autonomously to inhibit growth cone protrusion.

**Figure 8.**
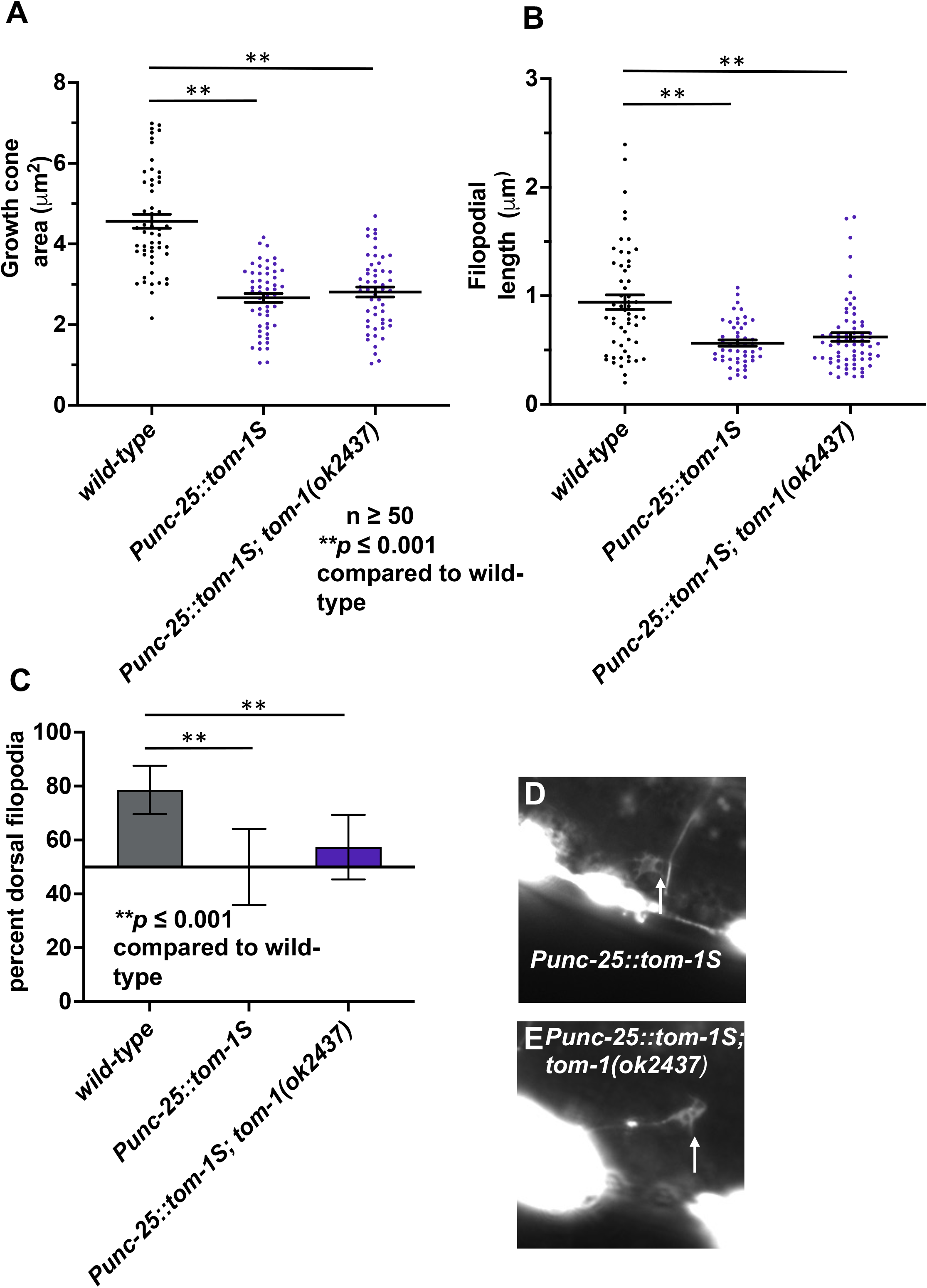
Growth cone analysis of transgenic expression of *tom-1 short*. (A, B) Quantification of VD growth cone area and filopodial length as described in Figure 4. Comparisons of significance indicated on previous figures are not shown here. Lines indicate significance of comparisons between genotypes. C) A graph showing the percent of dorsally-directed filopodial protrusions as described in Figure 4. (D-I) Fluorescence micrographs of mutant VD growth cones of indicated genotypes. The *gfp* fluorescence is from the *Punc-25::tom-1S::gfp* transgene. Arrows point to filopodial protrusions. Dorsal is up; anterior is left. The scale bar in (D) represents 5μm.

### unc-64/syntaxin regulates growth cone protrusion and genetically interacts with unc-5

Tomosyn inhibits vesicle fusion by interacting with the T-SNARE Syntaxin via the Tomosyn C-terminal R-SNARE domain, preventing syntaxin interaction with the vesicle SNARE VAMP (Fujita *et al*. 1998; Rizo and Rosenmund 2008). *unc-64* encodes for the *C. elegans* homolog of Syntaxin (Saifee *et al*. 1998). Complete loss of *unc-64* is lethal, but the hypomorphic *unc-64(md130)* allele is viable and displays slightly uncoordinated locomotion (Metz *et al*. 2007).

*unc-64(md130)* mutants displayed VD growth cones with reduced area and filopodial length compared to wild-type (Figure 9 A, B, and D). Furthermore, *unc-64(md130)* growth cones were unpolarized (Figure 9C). Finally, *unc-64(md130)* suppressed excessive growth cone protrusion in *unc-5* null and hypomorphic mutants (Figure 9 A, B, E, and F). *unc-64(md130)* displayed weak axon guidance defects alone, but double mutants of *unc-64; unc-5* displayed a synergistic enhancement in VD/DD axon guidance (Figure 3).

**Figure 9.**
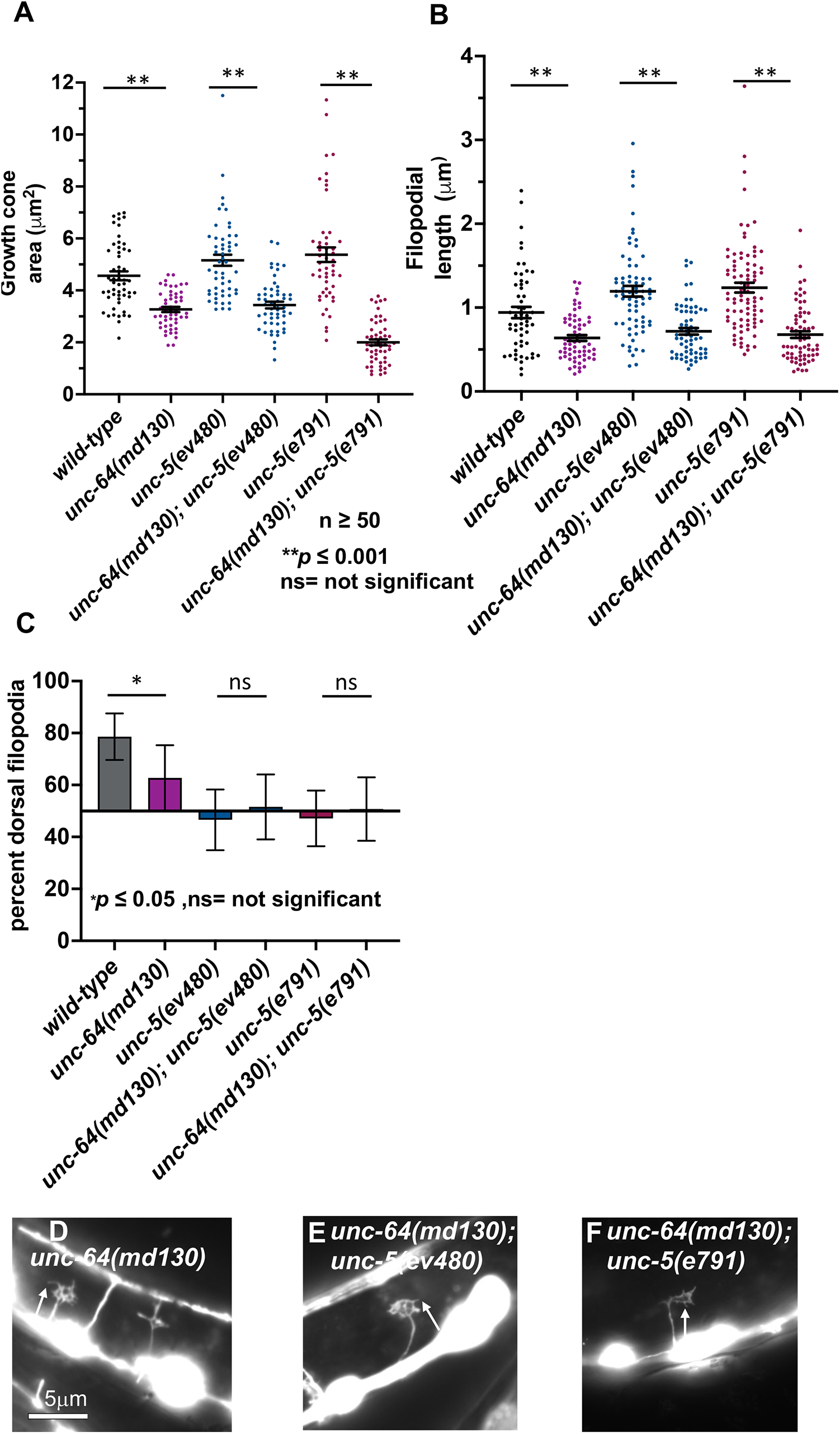
Growth cone analysis in *unc-64* and *unc-5* mutants. (A, B) Quantification of VD growth cone area and filopodial length as described in Figure 4. (C) A graph showing the percent of dorsally-directed filopodial protrusions in VD growth cones of different genotypes as described in Figure 4. (D-F) Fluorescence micrographs of mutant VD growth cones expressing *Punc-25::gfp*. Arrows point to filopodial protrusions. Dorsal is up; anterior is left. The scale bar in (D) represents 5μm.

These data indicate that UNC-64/Syntaxin is required for VD/DD growth cone protrusion, including growth cone area and filopodial length. They also suggest that in *unc-5* mutants, UNC-64/Syntaxin is overactive, resulting in excess growth cone protrusion.

## Discussion

Previous studies have shown that in the polarity/protrusion model of growth cone outgrowth, UNC-6/Netrin inhibits VD growth cone lamellipodial and filopodial protrusion via the UNC-5 receptor. UNC-6 also polarizes the VD growth cone via UNC-5, biasing filopodial protrusion to the dorsal direction of outgrowth. By inhibiting protrusion ventrally and stimulating protrusion dorsally (via the UNC-40 receptor), UNC-6 directs dorsal migration of the VD growth cone. Previous studies showed that UNC-5 inhibits growth cone protrusion using flavin monooxygenases (FMOs), which might oxidize and destabilize F-actin similar to MICAL. UNC-5 also inhibits protrusion by restricting entry of microtubule + ends into growth cones, which have a pro-protrusive effect. Results here suggest that UNC-5 inhibits protrusion via a third pathway involving UNC-64/Syntaxin and the Syntaxin inhibitor TOM-1/Tomosyn (Figure 10). UNC-64/Syntaxin was required for growth cone protrusion, and TOM-1/Tomosyn was an inhibitor of growth cone protrusion. Given the known roles of Syntaxin and Tomosyn in regulating vesicle fusion, these results are consistent with the idea that UNC-5 prevents vesicle fusion in the growth cone through TOM-1 inhibition of UNC-64/Syntaxin-mediated vesicle fusion. Prevention of vesicle fusion would restrict plasma membrane expansion and possibly delivery of pro-protrusive molecules (e.g. Arp2/3) and thus inhibit growth cone protrusion. *unc-64* and *tom-1* mutant VD growth cones were also unpolarized, indicating that regulation of vesicle fusion is required for establishing and/or maintaining growth cone polarity.

**Figure 10.**
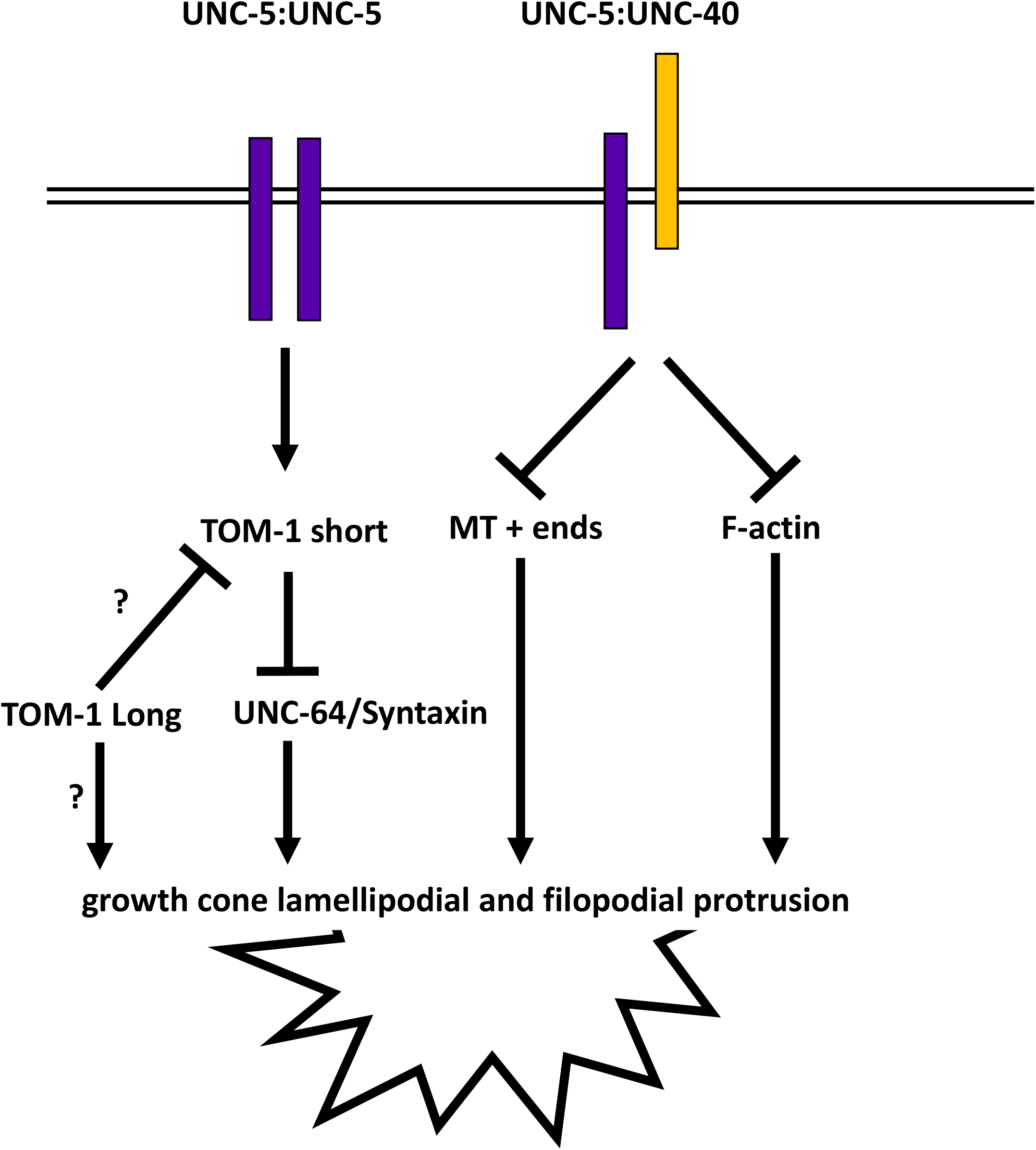
Genetic model of TOM-1 inhibiting growth cone protrusions by preventing vesicle fusion. Previous studies indicate that UNC-5 homodimers and UNC-5:UNC-40 heterodimers inhibit growth cone protrusion through the flavin monooxygenases (FMOs) and possible F-actin destabilization, and via UNC-33/CRMP and restriction of microtubule + end entry into growth cones. Studies here delineate a third pathway through which UNC-5 inhibits protrusion involving TOM-1/Tomosyn and the inhibition of vesicle fusion mediated by UNC-64/Syntaxin. Our results indicate that a short isoform of TOM-1 containing only the V-SNARE domain is the active isoform in inhibiting protrusion and possible interaction with UNC-64/Syntaxin. TOM-1 long isoforms might act in opposition to TOM-1 short and act in pro-protrusive manner. This might be a direct effect on protrusion, or possibly an auto-inhibitory effect on TOM-1 short.

Vesicle exocytosis is essential for controlling growth cone membrane dynamics and extensions by the fusion of plasmalemmal precursor vesicle at the plasma membrane of growth cones (Futerman and Banker 1996). Studies on rat cultured hippocampal neurons suggest that exocytosis is restricted to the peripheral region of growth cones which leads to the membrane addition and extension, do the action of Tomosyn in the growth cone “palm” (Sakisaka *et al*. 2004). *In vivo* results presented here are consistent with the idea that TOM-1 restricts vesicle fusion in the growth cone.

### *tom-1* encodes long and short isoforms with distinct functions

The *C. elegans* genome contains one *Tomosyn* gene, *tom-1*, which encodes N-terminal WD40 repeats and a C-terminal R-SNARE domain, similar to Tomosyn in other species. *tom-1* produced multiple isoforms, including a series of long isoforms containing the WD40 repeats and the R-SNARE domain, and a short isoform containing only the R-SNARE domain, produced by an alternative 5’ exon. Genetic results here indicate that the long isoforms have pro-protrusive roles in the growth cone, whereas the short isoform inhibits protrusion. The short-isoform-specific *tom-1(lq176)* mutant resembled the *tom-1* null, suggesting that the long isoforms cannot provide TOM-1 function in the absence of the short isoform. These data suggest that the TOM-1 short isoform is important for inhibiting growth cone protrusion, possibly through interaction with UNC-64/Syntaxin. Transgenic expression of TOM-1 short resulted in smaller growth cones with shorter filopodia, consist with this model. The long isoforms might have pro-protrusive roles independent of the short isoform. Alternately, the long isoforms might regulate the short isoform and inhibit its function (Figure 10).

### TOM-1 short isoform acts downstream of UNC-5 to inhibit the growth cone protrusion

Previous work showed that the UNC-6/Netrin receptors UNC-40 and UNC-5 regulate growth cone protrusion. UNC-40 stimulates protrusion whereas UNC-5 inhibits protrusion, and asymmetric distribution of protrusive activity across the growth cone results in directed growth cone migration away from UNC-6/Netrin (the Polarity/Protrusion model) (Gujar *et al*. 2018). We also showed that UNC-5 inhibits protrusion via the FMOs by possible actin destabilization (Gujar *et al*. 2017), and by preventing MT entry via UNC-33/CRMP (Gujar *et al*. 2017; Gujar *et al*. 2019).

Here we show that the TOM-1 short isoform is required for UNC-5 to inhibit VD growth cone protrusion. The *tom-1(ok2437)* null and the short-isoform-specific *tom-1(lq176)* both suppressed the inhibitory effect of MYR::UNC-5 on VD growth cone protrusion. While neither mutant alone had significantly increased growth protrusion expected of an inhibitory molecule, the effects might be masked by the other pathways downstream of UNC-5 (FMOs and UNC-33/CRMP). MYR::UNC-5 is a sensitized genetic background that revealed the effects on *tom-1* mutants on the VD growth cone. *tom-1(ok2437)* and *tom-1(lq176)* did not suppress the inhibitory effects on MYR::UNC-40, suggesting that TOM-1 might act specifically downstream of UNC-5 homodimers and not UNC-5:UNC-40 heterodimers. Finally, *tom-1(ok2437)* and *tom-1(lq176)* did not affect the large, overly protrusive growth cones of *unc-5* loss-of-function mutants, consistent with a role in inhibiting protrusion.

### TOM-1 long isoforms have a pro-protrusive role in the VD growth cone

In contrast to the *tom-1(ok2437)* null and *tom-1(lq176)* short isoform specific mutant, the long isoform-specific mutant *tom-1(nu468)* displayed VD growth cones with reduced area and filopodial length. *tom-1(nu468)* was required for the large, overly protrusive growth cones in *unc-5* null mutants, and did not suppress MYR::UNC-5. Together, these data indicate that TOM-1 long isoforms have a pro-protrusive role in the growth cone, the opposite of TOM-1 short. The TOM-1 long isoforms were required for excess protrusion in *unc-5* mutants, suggesting that in the absence of UNC-5, the TOM-1 long isoforms are overactive.

Previous neurophysiological studies indicate that the TOM-1 long isoforms are inhibitory to neurosecretion (Dybbs *et al*. 2005; Gracheva *et al*. 2006; McEwen *et al*. 2006; Gracheva *et al*. 2010; Burdina *et al*. 2011). Expression of the R-SNARE domain alone was insufficient to restore inhibition of synaptic transmission (Burdina *et al*. 2011), whereas experiments here show that expression of the TOM-1 short isoform inhibited VD growth cone protrusion. Possibly, the function of the long and short isoforms in vesicle fusion are cell-and context-specific. Indeed, in cultured superior cervical ganglion neurons, Tomosyn RNAi inhibited the evoked response (Baba *et al*. 2005), the opposite of what is expected of an inhibitor of vesicle fusion. In the VD growth cone, TOM-1 long might act as a true “friend to Syntaxin”, possibly inhibiting the function of TOM-1 short. It is also possible that TOM-1 long isoforms have a Syntaxin-independent stimulatory effect on growth cone protrusion.

### TOM-1 long and short isoforms are both required for VD growth cone polarity and VD/DD axon guidance

All *tom-1* mutants analyzed here displayed loss of dorsally-polarized filopodial protrusions on the growth cone. Transgenic expression of the TOM-1 short also resulted in VD growth cone polarity defects. This indicates that both TOM-1L and TOM-1S are required to establish and/or maintain VD growth cone polarity in a complex and likely dynamic manner. No genetic interaction analyzed here modified the polarity defect, so it is impossible to say if TOM-1 acts downstream of UNC-5 in growth cone polarity.

All *tom-1* alleles displayed weak but significant VD/DD axon guidance defects, and all of them synergistically enhanced the VD/DD axon guidance defects in double mutant combinations with *unc-5*. This suggests that both isoforms of *tom-1* are necessary to maintain proper axon guidance and again supports the hypothesis that TOM-1 acts downstream of UNC-5 signaling.

### UNC-64/Syntaxin is required for robust VD growth cone protrusion and polarity

Tomosyn is very well characterized as an inhibitor of vesicle fusion by blocking interaction of Syntaxin with the V-SNARE Synaptobrevin, including *C. elegans* TOM-1 in neurosecretion (Dybbs *et al*. 2005; Gracheva *et al*. 2006; McEwen *et al*. 2006; Gracheva *et al*. 2010; Burdina *et al*. 2011). It is possible that the effects of TOM-1 in growth cone protrusion are independent of vesicle fusion. However, hypomorphic *unc-64/Syntaxin* mutans displayed reduced VD growth cone area, shorter filopodial protrusions, and a loss of polarity of filopodial protrusions. This is similar to the long-isoform-specific *tom-1(nu468)* mutant and is consistent with a role of UNC-64/Syntaxin in promoting growth cone protrusion and polarity. This strongly indicates that vesicle fusion is required for robust VD growth cone protrusion, filopodial protrusion, and polarity of filopodial protrusion. The effects of TOM-1 on these processes are thus likely to be mediated through regulation of vesicle fusion.

### The UNC-6/Netrin UNC-5 inhibits VD growth cone protrusion via TOM-1/Tomosyn

In cultured rat hippocampal neurons, Tomosyn prevents vesicle fusion at the “palm” of the growth cone, directing vesicle fusion to the extending growth cone tip (Sakisaka *et al*. 2004). Evidence is presented here *in vivo* in *C. elegans* that TOM-1/Tomosyn might act similarly in the VD growth cone. Tom-1/Tomosyn might be active at the base of the VD growth cone, preventing ventral and lateral vesicle fusion and thus preventing ventral and lateral growth cone protrusion. We show that TOM-1 acts downstream of the UNC-6/Netrin receptor UNC-5 to inhibit protrusion. As UNC-5 also polarizes the growth cone, the activity of TOM-1 ventrally and laterally might be controlled by UNC-5. TOM-1 is also required to establish and/or maintain growth cone polarity of protrusion, suggesting the role of vesicle fusion in this process as well. In cultured rat hippocampal neurons, upon growth cone collapse, Tomosyn extends throughout the growth cone. This situation might be analogous to constitutively-active MYR::UNC-5, which might constitutively recruit TOM-1 throughout the entire growth cone, leading to inhibited protrusion. These results advance understanding of the role of UNC-6/Netrin receptor UNC-5 in growth cone morphology during outgrowth. They show that, in addition to the two pathways involving F-actin and microtubule + end entry, UNC-5 inhibits protrusion by preventing vesicle fusion in the growth cone using TOM-1/Tomosyn.

## Materials and Methods

### Genetic Methods

Experiments were performed at 20^0^C using standard *C. elegans* techniques (Brenner 1974). Mutations were used LGI: *tom-1(ok2437, nu468, lq176), wrdSi23* ; LGII: *juIs76* [*Punc-25*::*gfp*]; LGIII: *unc-64(md130)*; LGIV: *unc-5(e791, ev480* and, *e152)*, The presence of mutations was confirmed by phenotype and sequencing. Chromosomal locations not determined: *lqIs296[Punc-25::myr::unc-5::gfp]; lqIs128[Punc-25::myr::unc-40::gfp]; lqIs383 [Punc-25::tom-1short::gfp]*.

### Transgene construction

*Punc-25::tom-1 short* was made by amplifying the entire genomic region of the *tom-1B* short isoform by PCR, and placing this behind the *unc-25* promoter. The *tom-1B* coding region was sequenced to ensure no errors had been introduced by PCR. The sequence of this plasmid is available upon request.

### Cas9 genome editing to generate *tom-1B(lq176)*

CRISPR-Cas9 genome editing was used to precisely delete the entire intron 18 of *tom-1A*, between exon 18 and 19 (Figure 1). This removes the first exon of the *tom-1B* short isoform, which resides in intron 18, and leaves the coding potential of *tom-1* long isoforms unchanged. Synthetic guide RNAs were directed at the 5’ and 3’ ends of the intron:

sgRNA1 against *tom-1* short: 5’ CATCAATTTCCACAGAATGT 3’ sgRNA2 against *tom-1* short: 5’ TTACATGGCAAGTCAAACAG 3’

A mix of sgRNAs, a single stranded oligonucleotide repair template, and Cas9 enzyme was injected into the gonads of N2 animals, along with the *dpy-10(cn64)* co-CRISPR reagents (El Mouridi *et al*. 2017). Deletion of *tom-1* intron 18 was confirmed by PCR and sequencing. A single-stranded oligodeoxynucleotide was used as a repair template, which recoded the sgRNA region to maintain the same coding potential. The recoded sequence of *lq176* is: TTGAGAATTCTTCAAACTTCTACGATTTTCCCACACTCCGTCGAGATCGACGACCCA CTCTGCCAAAAGACCGCCTTCTCCGACCATGGACTCGGAGTCTATATGGCCTCCC AAACAGAGGTAAGATACTTTGTTTTATCATGAAAGTTA

The mutation was named *tom-1(lq176)*. Genome editing reagents were produced by InVivo Biosystems (Eugene, OR, USA).

### RNA-seq and analysis

Total RNA was isolated from three independent isolates of mixed-stage animals of the strain LE6194 (SSM1, SSM2, and SSM3) as previously described (Tamayo *et al*. 2013). The LE6194 genotype is *wrdSi23 I; juIs76 II* and is wild-type for the *tom-1* gene. Stranded poly-A RNA-seq libraries were constructed using the NEBNext® Ultra™ II Directional RNA Library Prep Kit for Illumina and subjected to paired-end 150 cycle sequencing on a Nextseq550. Reads were aligned to the *C. elegans* genome using HISAT2 with default settings (version 2.1.0) (Kim *et al*. 2019). The resulting BAM files were analyzed in the Integrated Genome Viewer (Robinson *et al*. 2011; Thorvaldsdottir *et al*. 2012) from which the Sashimi plot for SSM1 in Figure 1B was generated. The following numbers of paired reads for each sample mapped to the genome: SSM1, 46,067,340; SSM2, 52,240,104; SSM3, 55,063,010. Sequences are available in the Sequence Read Archive, BioProject number PRJNA847250.

### Quantification of axon guidance defects

VD/DD neurons were visualized with *Punc-25*::*gfp* transgene, *juIs76* (Jin *et al*. 1999), which is expressed in GABAergic motor neurons including 13VDs and 6DDs. Axon guidance defects were scored as previously described (Mahadik and Lundquist 2022). In wild type, an average of 16 of the 19 commissures of VD/DD axons are distinguishable, as axons can be present in a fascicle and thus cannot be resolved. A total of 100 animals were scored (1600 total commissural processes). Total axon guidance defects were calculated by counting all the axons which failed to reach the dorsal nerve cord, wandered at an angle of 45^0^ or greater, crossed over other processes, and displayed ectopic branching. Significance difference between two genotypes was determined by using Fisher’s exact test.

### Growth cone imaging and quantification

VD growth cones were imaged as previously described (Norris and Lundquist 2011; Norris *et al*. 2014; Gujar *et al*. 2017; Gujar *et al*. 2018; Gujar *et al*. 2019; Mahadik and Lundquist 2022). L1 larval animals were harvested 16-hour post-hatching at 20^0^C and placed on a 2% agarose pad with 5mM sodium azide in M9 buffer. Some genotypes were slower to develop than others so the 16-hour timepoint was adjusted for each genotype. Growth cones were imaged with Qimaging Retiga EXi camera on a Leica DM5500 microscope at 100x magnififcation. Projections less than 0.5um in width were scored as filopodia. Growth cone area and filopodial length were quantified using ImageJ software. Quantification was done as described previously (Norris and Lundquist 2011; Norris *et al*. 2014; Gujar *et al*. 2017; Gujar *et al*. 2018; Gujar *et al*. 2019; Mahadik and Lundquist 2022). Significance of difference between two genotypes was determined by two-sided *t*-test with unequal variance.

Polarity of growth filopodial protrusions was determined as previously described (Norris and Lundquist 2011; Norris *et al*. 2014; Gujar *et al*. 2017; Gujar *et al*. 2018; Gujar *et al*. 2019; Mahadik and Lundquist 2022). Growth cone images were divided into two halves, dorsal and ventral, with respect to the ventral nerve cord. The number of filopodia in each half was counted. The proportion of dorsal filipodia was determined by the number of dorsal filopodia divided by total number of filopodia. Significance of difference between two genotypes was determined by Fisher’s exact test.

## Acknowledgments

The authors thank B. Ackley, Lundquist and Ackley lab members, and J. Richmond for helpful discussion. Some *C. elegans* strains were provided by the CGC, which is funded by NIH Office of Research Infrastructure Programs (P40 OD010440).

## Funding

National Institutes of Health R03 NS114554 to E.A.L.

National Institutes of Health R56 NS095682 to E.A.L.

National Institutes of Health P20 GM103638 (S. Lunte PI, E.A.L. coI).

## Core Labs

Next generation sequencing was performed by he KU Genome Sequencing Core Laboratory, part of the *Center for Molecular Analysis of Disease Pathways* (National Institutes of Health P20 GM103638 (S. Lunte PI, E.A.L. coI)).

